# Vertebral level specific modulation of paraspinal muscle activity based on vestibular signals during walking

**DOI:** 10.1101/2023.10.18.562858

**Authors:** Yiyuan C. Li, Sjoerd M. Bruijn, Koen K. Lemaire, Simon Brumagne, Jaap H. van Dieën

**Affiliations:** Department of Human Movement Sciences, Faculty of Behavioral and Movement Sciences, Vrije Universiteit Amsterdam, Amsterdam Movement Sciences, Amsterdam, The Netherlands; Department of Rehabilitation Sciences, KU Leuven, Leuven, Belgium

**Keywords:** electrical vestibular stimulation, paraspinal muscles, balance control, gait stability

## Abstract

Evoking muscle responses by electrical vestibular stimulation (EVS) may help to understand the contribution of the vestibular system to postural control. Although paraspinal muscles play a role in postural stability, the vestibulo-muscular coupling of these muscles during walking has rarely been studied. This study aimed to investigate how vestibular signals affect paraspinal muscle activity at different vertebral levels during walking with preferred and narrow step width. Sixteen healthy participants were recruited. Participants walked on a treadmill for 8 minutes at 78 steps/min and 2.8 km/h, at two different step width, either with or without EVS. Bipolar electromyography was recorded bilaterally from the paraspinal muscles at eight vertebral levels from cervical to lumbar. Coherence, gain, and delay of EVS and EMG responses were determined. Significant EVS-EMG coupling (p<0.01) was found at ipsilateral and/or contralateral heel strikes. This coupling was mirrored between left and right relative to the midline of the trunk and between the higher and lower vertebral levels, i.e., a peak occurred at ipsilateral heel strike at lower levels, whereas it occurred at contralateral heel strike at higher levels. EVS-EMG coupling only partially coincided with peak muscle activity. EVS-EMG coherence slightly, but not significantly, increased when walking with narrow steps. No significant differences were found in gain and phase between the vertebral levels or step width conditions. In summary, vertebral level specific modulation of paraspinal muscle activity based on vestibular signals might allow a fast, synchronized, and spatially co-ordinated response along the trunk during walking.

**Key Points Summary:**

- Mediolateral stabilization of gait requires an estimate of the state of the body, which is affected by vestibular afference.
- During gait, the heavy trunk segment is controlled by phasic paraspinal muscle activity and in rodents the medial and lateral vestibulospinal tracts activate these muscles.
- To gain insight in vestibulospinal connections in humans and their role in gait, we recorded paraspinal surface EMG of cervical to lumbar paraspinal muscles, and characterized coherence, gain and delay between EMG and electrical vestibular stimulation, during slow walking.
- Vestibular stimulation caused phasic, vertebral level specific modulation of paraspinal muscle activity at delays of around 40 milliseconds, which was mirrored between left, lower and right, upper vertebral levels.
- Our results indicate that vestibular afference causes fast, synchronized, and spatially co-ordinated responses of the paraspinal muscles along the trunk, that simultaneously contribute to stabilizing the centre of mass trajectory and to keeping the head upright.

## Introduction

Stable walking, defined as “gait that does not lead to falls” (Bruijn *et al*., 2013), is an effortless and seemingly simple task for healthy young adults. However, with the presence of disorder (e.g., low back pain, bilateral vestibular loss, etc.), aging, or external perturbations, the difficulty of the task becomes manifest (Mazaheri *et al*., 2013; Sprenger *et al*., 2017). To maintain stability, the centre of mass should be controlled relative to the base of support. Three mechanisms can contribute to the control of the centre of mass: modulation of foot placement, shifting the centre of pressure through ankle moments, and changing angular momentum such as through trunk rotation (known as counter-rotation) (Hof, 2007).

These mechanisms for stability are believed to be controlled using the error between efference copies and the actual state detected by sensory systems. The vestibular system, for instance, which encodes head motion, has been shown to contribute to gait stabilization (van Schooten *et al*., 2011; Cullen, 2012; Magnani *et al*., 2021). It has been reported that the medial vestibulospinal tract, commencing in the medial vestibular nucleus and then bilaterally extending through the cervical spinal cord into the medial longitudinal fasciculus, is involved in head stabilization (Wilson & Maeda, 1974; Fukushima *et al*., 1979; Goldberg & Cullen, 2011; Forbes *et al*., 2020). Another descending brainstem pathway involved in stabilization is the lateral vestibulospinal tract, which originates in the lateral vestibular nucleus and projects ipsilaterally to axial motoneurons at all spinal levels (Grillner & Hongo, 1972; Fukushima *et al*., 1979; Basaldella *et al*., 2015). This pathway is required for the preservation of upright posture and for responding to postural perturbations (Markham, 1987; Dichgans & Diener, 1989; Murray *et al*., 2018). The vestibulospinal reflex has been investigated experimentally by applying electrical vestibular stimulation (EVS). This stimulation activates the vestibular afferent nerve fibres without head movements and produces an illusion of moving the head (Goldberg *et al*., 1982; Day, 1999; Kwan *et al*., 2019). Consequently, muscle responses can be detected in balance related muscles while standing and walking (Ali *et al*., 2003; Dakin *et al*., 2010; Guillaud *et al*., 2020; Magnani *et al*., 2021). Studies have revealed that these muscle responses are frequency, task, and muscle specific (Dakin *et al*., 2010; Forbes *et al*., 2013).

While the responses of lower limb muscles to vestibular stimulation during walking have been extensively investigated (Forbes *et al*., 2013; Magnani *et al*., 2021), little attention has been given to the trunk muscle responses to EVS, despite the fact that the upper body represents approximately 50% of the total body mass (de Leva, 1996). In the study of gait, most attention has been devoted to the control of the legs, whereas body parts above the hips are commonly considered as a single rigid head-arm-trunk segment. However, in terms of maintaining stability, the trunk may play a significant role, as even a tiny deviation from the perfect upright position can produce a substantial destabilizing moment (Bruijn & van Dieen, 2018). Additionally, the trunk may be crucial when employing the counter-rotation mechanism for stabilization (Hof, 2007).

A narrow step width, which reduces the base of support, poses greater challenges to stability and, as such, increases the demands on postural control (Arvin *et al*., 2016; Magnani *et al*., 2023). A previous study demonstrated that vestibular contributions are modulated by the stabilization demands of walking (Magnani *et al*., 2021). This study showed an increased coherence between EVS and ground reaction force during walking with narrow steps but decreased EVS-ground reaction force coherence during walking with wider steps. Though the coherence between EVS and EMG of lower limb muscles remained unchanged or decreased, the EVS-EMG coherence increased in the paraspinal muscles at the lumbar region when walking with a narrow step width (Magnani *et al*., 2021). Overall, these findings indicate the importance of coupling between vestibular signals and paraspinal muscles in controlling stability during walking. However, while this previous study did look at the paraspinal muscles, it only studied the paraspinal muscles at one vertebral level of the trunk. It is therefore unknown if, and how, paraspinal muscles at other vertebral levels are involved in the control of gait stability.

Our research question was: If and how do paraspinal muscles at different vertebral levels respond to electrical vestibular stimulation throughout the gait cycle? Furthermore, we tested if these vestibular contributions varied with step width. Based on previous studies (Forbes *et al*., 2013; Magnani *et al*., 2021), we hypothesized that: 1) paraspinal muscle show activity coherent with EVS at specific phases of the gait cycle, specifically at heel strike; 2) the EVS-evoked muscles responses increase when walking with narrow step widths. To test our hypotheses, we characterized the coherence, gain and delay of EMG-electrical vestibular stimulation response in paraspinal muscle during walking with both preferred step width and narrow step width. Paraspinal muscles from cervical to lumbar vertebral levels were studied.

## Methods

### Ethical Approval

The study conformed to the standards set by the Declaration of Helsinki, except for registration in a database. Procedures had been approved by the VU Amsterdam Research Ethics Committee VCWE-2022-126.

### Participants

Sixteen healthy participants (7 male, 9 female) between the ages of 19 and 29 years were recruited (Age: 23.5 ± 3.4 years, height: 1.71 ± 0.07 m, weight: 64.5 ± 13.2 kg). Exclusion criteria for participation included any diagnosed orthopaedic or neurological disorders, or the use of medications that can cause dizziness. Participants were instructed not to engage in intense physical exercise and to refrain from alcohol consumption 24 hours prior to the experiment. Two participants were excluded during data analysis due to technical problems in the signal synchronization between devices. The participants agreed to participate in the study by signing an informed consent form.

### Electrical Vestibular Stimulation

Continuous EVS was applied in a bipolar configuration. The EVS was delivered as an analogue signal via a digital-to-analogue converter (National Instruments Corp., Austin, USA) to an isolated constant-current stimulator (BIOPAC System Inc., Goleta, USA) that was connected to carbon rubber electrodes (9 cm²). The two electrodes were coated with electrode gel (SonoGel, Bad Camberg, Germany) and placed over the mastoid processes, fixed tightly with hypoallergenic tapes and an elastic head band (Figure 1A).

**Figure 1.**
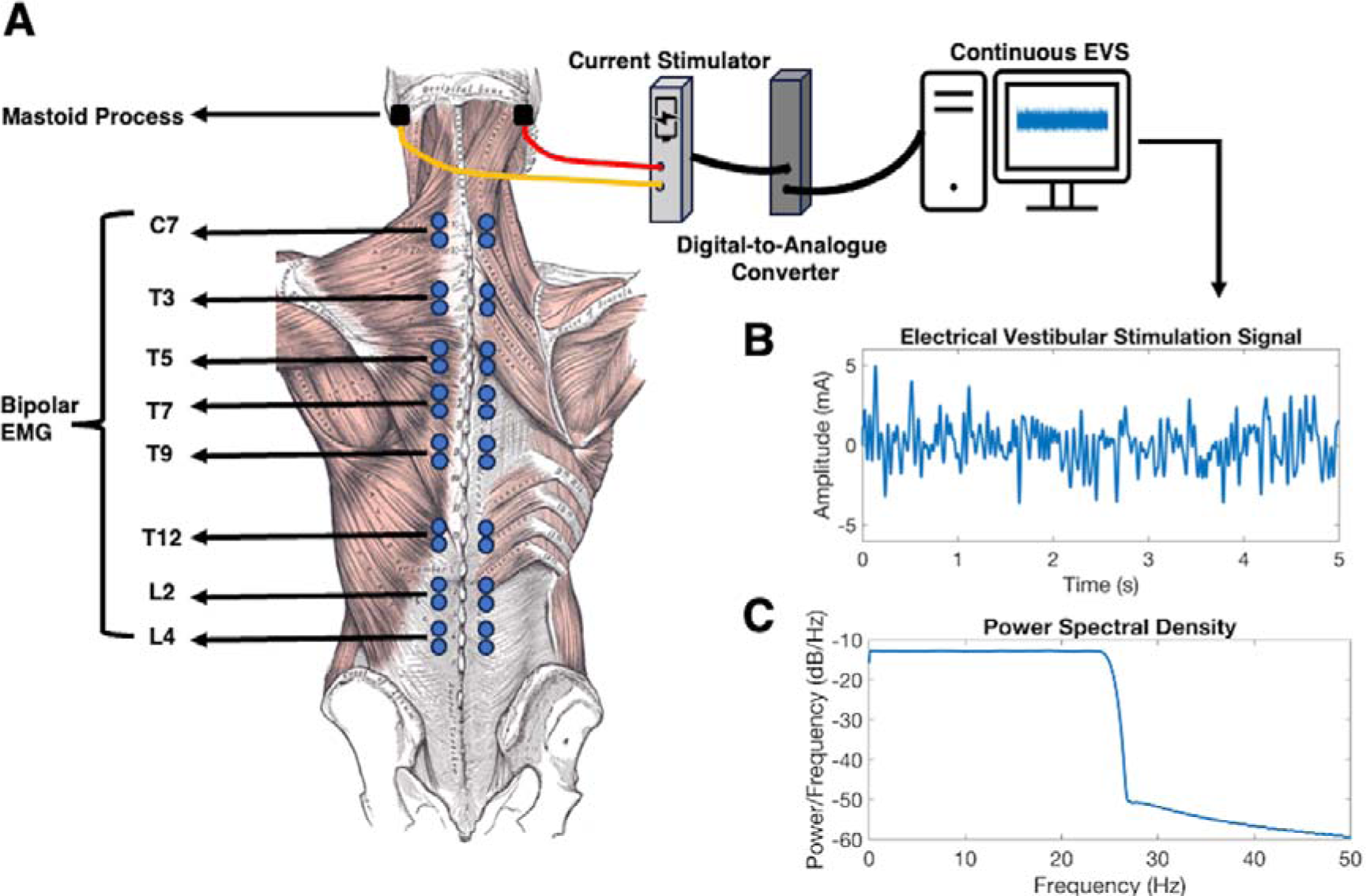
Experimental Setup. (A) The locations of the EVS electrodes (black squares) over the mastoid process, and paired EMG electrodes (blue dots) over the paraspinal muscles at eight vertebral levels from top to bottom: C7, T3, T5, T7, T9, T12, L2 and L4. EMG electrodes were placed 2 to 3 cm lateral to the mid-sagittal line defined by the spinous processes. (B) Time series of the electrical vestibular stimulation signal, zoomed in to a window of 5 seconds. (C) The power spectral density of the electrical vestibular stimulation signal.

All participants were exposed to the same zero-mean stochastic EVS signal, which was a white noise generated in Matlab (2019a, The MathWorks, Natick, US) with a bandwidth from 0 to 25 Hz, peak amplitude of ± 5.0 mA, root mean square ofll∼ll1.2 mA, and duration of 8 minutes (Figure 1B & 1C). These signal characteristics were identical to those used previously (Magnani *et al*., 2021; Magnani *et al*., 2023), in which measurable responses to EVS in muscle activity, ground reaction force and foot placement were reported, demonstrating the effectiveness of this EVS protocol.

### Electromyography and Ground Reaction Force Measurement

Bipolar surface electromyography (EMG) was recorded by a 64-channel REFA amplifier (TMSi, Oldenzaal, The Netherlands; CMRR: > 90 dB; Input impedance: > 100 MΩ; Gain: 26.55), bilaterally from the paraspinal muscles at 2048 samples/s using disposable self-adhesive Ag/AgCl surface electrodes (Ambu blue sensor, Ballerup, Denmark). EMG was recorded from paraspinal muscles at eight vertebral levels, including the seventh cervical vertebra (C7), the third, fifth, seventh, nineth and twelfth thoracic vertebra (T3, T5, T7, T9, T12) and the second and fourth lumbar vertebra (L2, L4). After shaving, lightly abrading, and cleaning the skin with alcohol, electrodes were placed 2 to 3 cm lateral to the mid-sagittal line, defined by the spinous processes (Figure 1A). A reference electrode was placed over the bony part of the acromion process.

Ground reaction forces (GRF) were measured at a sampling rate of 1000 Hz by force plates, which were built-in the split-belt treadmill (ForceLink b.v., Culemborg, the Netherlands). The force plate data were used to identify heel-strike. Kinematic data were recorded using a 3D motion capture system (Northern Digital Inc, Waterloo Ont, Canada), sampling at 50 samples/s, to calculate the step widths and the variability of head and thorax rotation.

### Protocol

Prior to the start of the experiment, participants were asked to walk on the dual-belt treadmill at a speed of 2.8 km/h without EVS for 3 minutes to measure their preferred step widths (PSW). For familiarization, participants were exposed to vestibular stimulation in a three-minutes-walk at 2.8 km/h. In addition, they were instructed to time their steps to the beat of a metronome at 78 steps/min.

For the experiment, participants walked on the dual-belt treadmill at 2.8 km/h with a cadence of 78 steps/min for 8 minutes. Walking speed and cadence were chosen to replicate the conditions of Magnani et al. (2021). The conditions were defined by the presence of EVS (EVS and no-EVS), by the step width (preferred step width: PSW and narrow step width: NSW) and by head orientation (facing forward and facing sideways). Participants were instructed to face forward in all conditions except in head sideways condition. Data from head sideways condition were not used in the current study. NSW was defined as 50% of PSW. To impose step width, two lines were projected on the treadmill, symmetrical to its midline. Participants were instructed to align the middle of their shoes with these lines and to check the line with peripheral vision (Arvin *et al*., 2016). The order of the conditions was randomized for each participant. Throughout the conditions, participants received repeated verbal instruction to maintain their cadence and step width. Before each condition, participants were given sufficient time to rest, followed by 2 min walking without vestibular stimulation before data collection, to avoid acclimatization and/or habituation effects.

### Signal Analysis

The electromyography, force plate and vestibular stimulation signals were first synchronized based on the trigger signals. Subsequently, EMG signals (2048 Hz) were resampled to 1000 Hz, aligned with the sampling rate of the force plate and EVS signals. Time instances of heel strikes were calculated from force plate data from the characteristic “butterfly pattern” of the centre of pressure (Roerdink *et al*., 2008). To test if, and how well participants were able to execute the task of controlling step widths and cadence, the mean step width of each participant was calculated as the mediolateral distance between the second toe tip markers during heel strikes. The cadence was calculated from the time between subsequent heel strikes of the same leg.

To investigate the effects of EVS on stability control, the variability of head and thorax rotation in the global coordinate system was calculated. The global coordinate system was defined as positive y-axis forward, positive x laterally to the right, positive z-axis vertically upward. Time series of thorax and head orientation were calculated using Euler decomposition (y-x-z sequence). The variability of rotation was calculated as the standard deviation of the orientation angle over gait cycles at each normalized time point, subsequently averaged across the gait cycle for each participant.

The vestibular stimulation caused artifacts in the EMG signals, particularly when recording from the cervical and upper thoracic region (Ali *et al*., 2003). To assess how to remove these artifacts, we compared the power spectral density of filtered EMG signals and the EVS-EMG coherence after filtering with different cut-off frequencies (30, 40, 50, 60, 70, 80, 90, 100 and 110 Hz) (Forbes et al., 2013). In the end, the bipolar EMG signals were high-pass filtered with a 6th order, Butterworth high-pass filter with a cut-off frequency of 100 Hz, in line with previous literature (Blouin et al., 2011; Forbes et al., 2013). Filtered EMG signals were rectified and normalized to the maximum amplitude of EMG from the averaged gait cycle in the No-EVS-PSW condition for each vertebral level and participant.

Estimates of coherence, gain and phase were based on the continuous Morlet wavelet decomposition (Blouin *et al*., 2011). Before calculating the auto- and cross-spectra, EVS and normalized EMG signals were first sectioned into strides based on the right heel strikes. To avoid distortion in the coherence and gain calculation at either the beginning or the end of the signals, each stride was padded with data from the previous (50%) and subsequent (50%) strides. Time-frequency coherence, *Coh(τ, f),* was computed to measure the linear relationship between the EVS and EMG signals across the frequencies considered as:

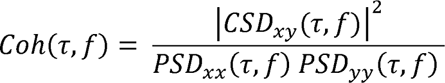

Where τ and *f* denote the stride time and frequency, *CSD_XY_* (τ, *f*) refers to the time-frequency cross-spectra between EVS and normalized EMG. *PSD_XX_(τ, f)* and *PSD_YY_ (τ, f*) are the time-frequency auto-spectra of EVS and EMG, respectively. To account for the variability of stride duration, time was normalized by resampling the *CSD_XY_*, *PSD_XX_ (τ, f)* and *PSD_YY_ (τ, f)* to 1000 samples per stride cycle. Coherence, which is unitless, ranges from 0 to 1, with 0 indicating independence and 1 indicating a perfect linear relationship between the two signals. Under the hypothesis that the stimulation signal and output signal are independent, the distribution of coherence estimate can be written as (Zhan *et al*., 2006):

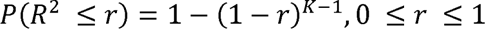

where *P(R*^2^ ≤ *r*) denotes the probability of coherence (*R*_2_) less than the detection threshold (*r*) and *K* is the number of strides for each participant. For a 99% confidence interval, corresponding to *p*ll<ll0.01, the threshold of coherence estimate is 0.015 with *K* as 300 strides, which was the minimum number of strides across all subjects.

To study the magnitude and timing of the output EMG relative to the input EVS, gain and phase estimates were calculated from the frequency response function (FRF), defined as:

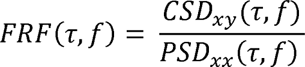

Where *FRF(τ, f)* indicates the time-frequency frequency response function. The FRF is a complex function, the modulus of which represents the gain (1/milliampere, mA^-1^), and the angle indicates the phase estimate (radian, rad). Since the gain and phase are not interpretable when the coherence between input and output is weak, we only used the phase and gain where the coherence reached its peak. The peak time-frequency coherence was defined as follows: first, the non-significant coherence estimates were removed for each participant and each vertebral level. The peak coherence was defined as the range of frequencies and gait phases where the coherence was larger than 99% of the remaining (significant) coherence estimates. Time-frequency phase estimates were then exacted from the region of peak coherence and transformed into time lags (millisecond, ms). Subsequently, the gain and delay were defined as the median of the gains and time lags across the frequencies and gait phases at peak coherence for each vertebral level and each participant. Gain and delay estimates from the left and right recording sites at the same vertebral level were averaged.

### Statistical Analysis

For the behavioural measures (i.e. step width, cadence, variability of head and thorax rotations), repeated-measures ANOVAs with two factors: Step width (preferred step width and narrow step width) × the Presence of EVS (with EVS and without EVS) were performed. To test the difference in muscle activity, repeated-measures analysis of ANOVAs with two factors: Step width (preferred step width and narrow step width) × the Presence of EVS (with EVS and without EVS) was performed via one-dimensional Statistical Parametric Mapping ((Pataky *et al*., 2016)). To identify the most prominent effects of step width and the electrical vestibular stimulation to muscle activity, significance was set as *p* value < 0.01. As the coherence ranges from 0 to 1, the modified Fisher-Z transform was applied on coherence before statistical analysis. For each participant, coherence exceeding 0.015, (i.e., pll<ll0.01), was defined as significant. To identify whether the paraspinal muscle responses significantly differed between step widths we performed cluster-based permutation tests (paired t-tests, 2000 permutations) on coherence in two step width conditions (Maris *et al*., 2007). To identify whether there was habituation to the effects of EVS, we performed cluster-based permutation tests (paired t-tests, 2000 permutations) on coherence in the first 50 gait cycles versus the last 50 gait cycles in both step width conditions. To test the differences in delays and gains, repeated-measures analysis of ANOVAs with two factors: Vertebral level (eight vertebral levels) × Step width (preferred step width and narrow step width) was performed. The assumption of sphericity for repeated-measures ANOVAs was verified by Mauchly’s Test (*p* >ll0.05). Greenhouse-Geisser correction was applied when the assumption of sphericity was rejected. A Bonferroni correction was applied for the post hoc tests. Significance was set at *p* < 0.05 for the statistical tests. Statistical analyses were performed in MATLAB (2019a, The MathWorks, Natick, US).

## Results

### Effects of EVS and Step Width on Gait Parameters

Although step width during walking with narrow step width (NSW) was not halved when compared to walking with preferred step width (PSW), it was significantly smaller than during walking with preferred step width (N = 14, F (0.520, 6.760) = 131.600, *p* < 0.001) (Figure 2A). This difference was observed both when walking with EVS (N = 14, *p* < 0.001) and without EVS (N = 14, *p* < 0.001), as confirmed by post hoc testing. EVS evoked a significant increase in step width (N = 14, F (0.520, 6.760) = 8.176, *p* = 0.013) when walking with preferred step width (N = 14, *p* = 0.038) and with narrow step width (N = 14, *p* = 0.016). No significant interaction between Step width and the Presence of EVS was found (N =14, F (0.520, 6.760) = 0.112, *p* = 0.742).

**Figure 2.**
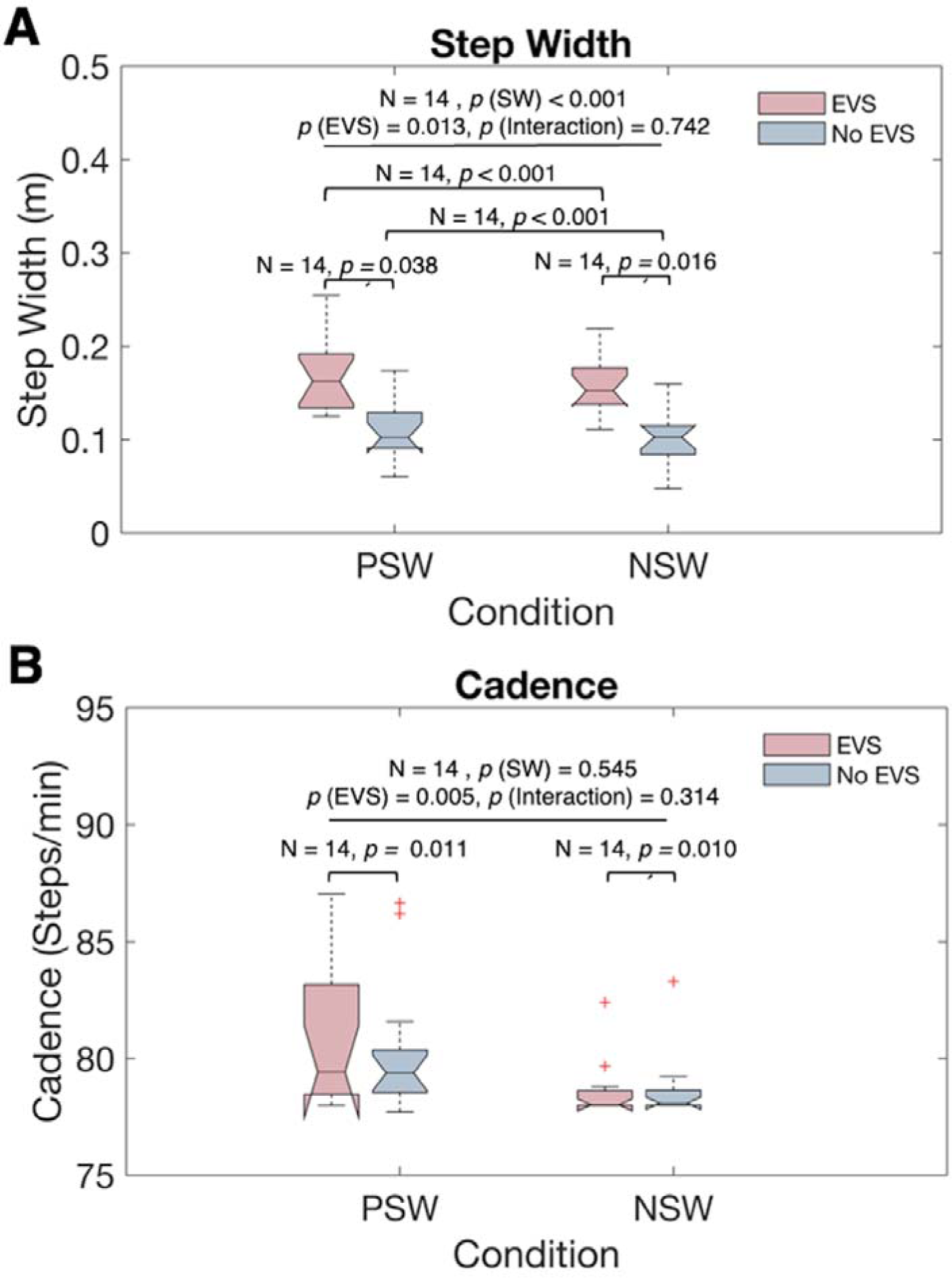
(A) Step Width and (B) Cadence. The X-axis indicate the step width conditions. The Y-axis represents the magnitude of step width (m) in panel A and cadence (steps/min) in panel B during walking with EVS (red box) and without EVS (blue box). Within each box, the central mark indicates the median, and the bottom and top edges indicate the 25th and 75th percentiles, respectively. The red ‘+’ symbols represent the outliers, and the whiskers extend to the maximum and minimum values, excluding the outliers

When walking without EVS, participants maintained their cadence as instructed (78 steps/min) with preferred step width (78.568 ± 1.202 steps/min) and narrow step width (78.647 ± 1.400 steps/min). However, in both conditions, EVS evoked a significant increase in cadence (N = 14, F (0.592, 7.690) = 11.043, *p* = 0.005) (Figure 2B). Post hoc testing indicated that the difference was significant both when walking with preferred step width (N = 14, *p* = 0.011) and with narrow step width (N = 14, *p* = 0.010). Step width (N =14, F (0.592, 7.690) = 0.385, *p* = 0.545) and the Step width × the Presence of EVS interaction (N =14, F (0.592, 7.690) = 1.438, *p* = 0.314) showed no significant effect on cadence.

### Effects of EVS and Step Width on Posture Control

Step width and the Presence of EVS showed a significant interaction effect on the head rotation variability (N = 14, F (0.849, 11.034) = 6.127, *p* = 0.028), while step width had a significant main effect on the head rotation variability in roll as well (N =14, F (0.849, 11.034) = 7.724, *p* = 0.016), but no significant effect was evoked by EVS (N =14, F (0.849, 11.034) = 2.712, *p* = 0.124) (Figure 3A). Post hoc testing indicated that when walking without EVS, narrow step width caused a significant decline in head rotation variability (1.236 ± 0.362 degree) compared to preferred step width (1.488 ± 0.532 degree) (N = 14, *p* = 0.008). In addition, a significant decrease in head rotation variability in roll was evoked by EVS when walking with preferred step width (N = 14, *p* = 0.014).

**Figure 3.**
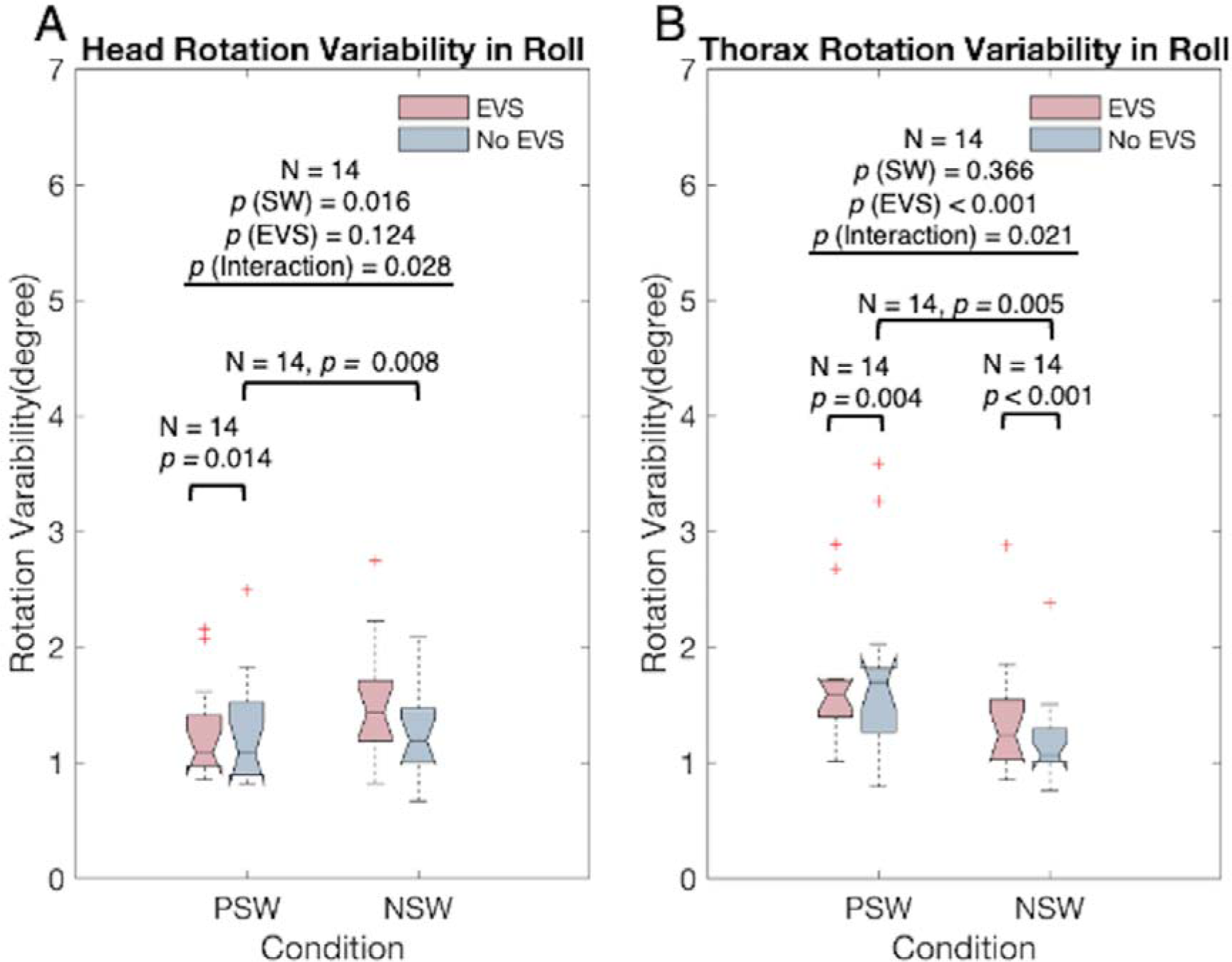
Variability of Head and Thorax Rotation in Two Step Width Conditions. The X-axis indicate the step width conditions. The Y-axis represents the variability of head rotation (degree) in panel A and thorax rotation (degree) in panel B during walking with EVS (red box) and without EVS (blue box).Within each box, the centre mark indicates the median, and the bottom and top edges indicate the 25th and 75th percentiles, respectively. The whiskers extend to the maximum and minimum values, excluding the outliers.

In contrast, no significant effect of step width on thorax rotation variability in roll was found (N = 14, F (0.481, 6.256) = 0.877, *p* = 0.366) (Figure 3B). However, the presence of EVS (N = 14, F (0.481, 6.256) = 19.579, *p* < 0.001) and Step width × Presence of EVS interaction (N = 14, F (0.481, 6.256) = 6.948, *p* = 0.021) showed significant effects on thorax rotation variability in roll. Based on post hoc testing, the exposure to EVS evoked a significant increase in thorax rotation variability in roll both when walking with preferred step width (N = 14, *p* = 0.004) and with narrow step width (N = 14, *p* < 0.001). When walking without EVS, thorax rotation variability was significantly larger in walking with preferred step width than narrow step width (N = 14, *p* = 0.005).

### Paraspinal Muscles Activity During Walking in Two Step Widths Conditions

Normalized muscle activity recorded from the paraspinal muscles at eight vertebral levels (C7, T3, T5, T7, T9, T12, L2, L4, bilaterally) during walking with and without electrical vestibular stimulation (EVS) with preferred step widths (PSW) and narrow step widths (NSW) is illustrated in Figure 4. At higher vertebral levels, including C7, T3, T5, T7 and T9, peaks occurred just after the contralateral heel strikes. At lower vertebral levels (T12, L2 and L4), muscle activity demonstrated two peaks occurring just after both ipsilateral and contralateral heel strikes.

**Figure 4.**
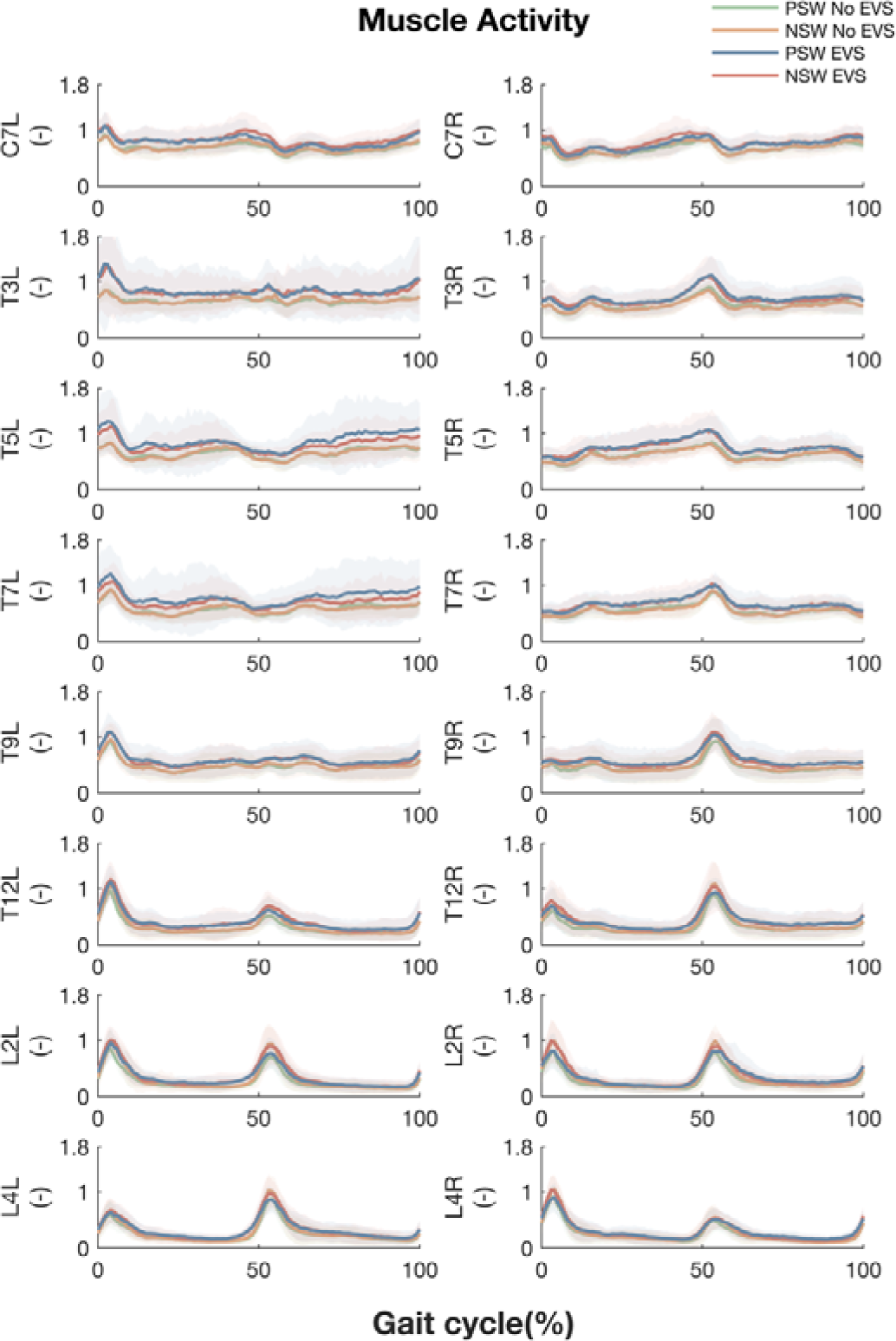
Paraspinal Muscle Activity Throughout the Gait Cycle. Averaged EMG linear envelopes of paraspinal muscles at all investigated vertebral levels during walking with EVS at preferred step width without EVS (Green), preferred step width with EVS (Blue), narrow step width without EVS (Orange) and narrow step width with EVS (Red). The X-axis indicates the time (normalized to gait cycle, starting at the right heel strike). The Y-axis presents the amplitude of the normalized EMG. The lines and shadowed areas represent the means and standard deviations across participants, respectively. The left and right columns depict recorded sites at the left and right side relative to the midline of the trunk. From top to bottom rows, the EMG of paraspinal muscles at eight vertebral levels are displayed as: C7, T3, T5, T7, T9, T12, L2 and L4.

Comparing the muscle activity in walking, the One-dimensional Statistical Parametric Mapping based repeated-measures ANOVA with two factors: Step width (preferred step width and narrow step width) × the Presence of EVS (with EVS and without EVS, featured supra-threshold clusters for the main effect ‘the Presence of EVS’ at the lower vertebral levels around ipsilateral heel strikes. For instance, at the left side of T12 at 47% gait cycles (*p* = 0.010), at the right side of T12 from 96 to 98% (*p* = 0.008), at the left side of L2 from 44 to 47% (*p* = 0.005), etc (Table 1). And there was a smaller region with a significant effect of ‘Step width’ at the right L2 and L4 vertebral levels, e.g., at around 7% of gait cycles (L2: *p* = 0.009; L4: *p* = 0.010). No significant interactions of EVS and step width on muscle activity were found.

**Table 1.**
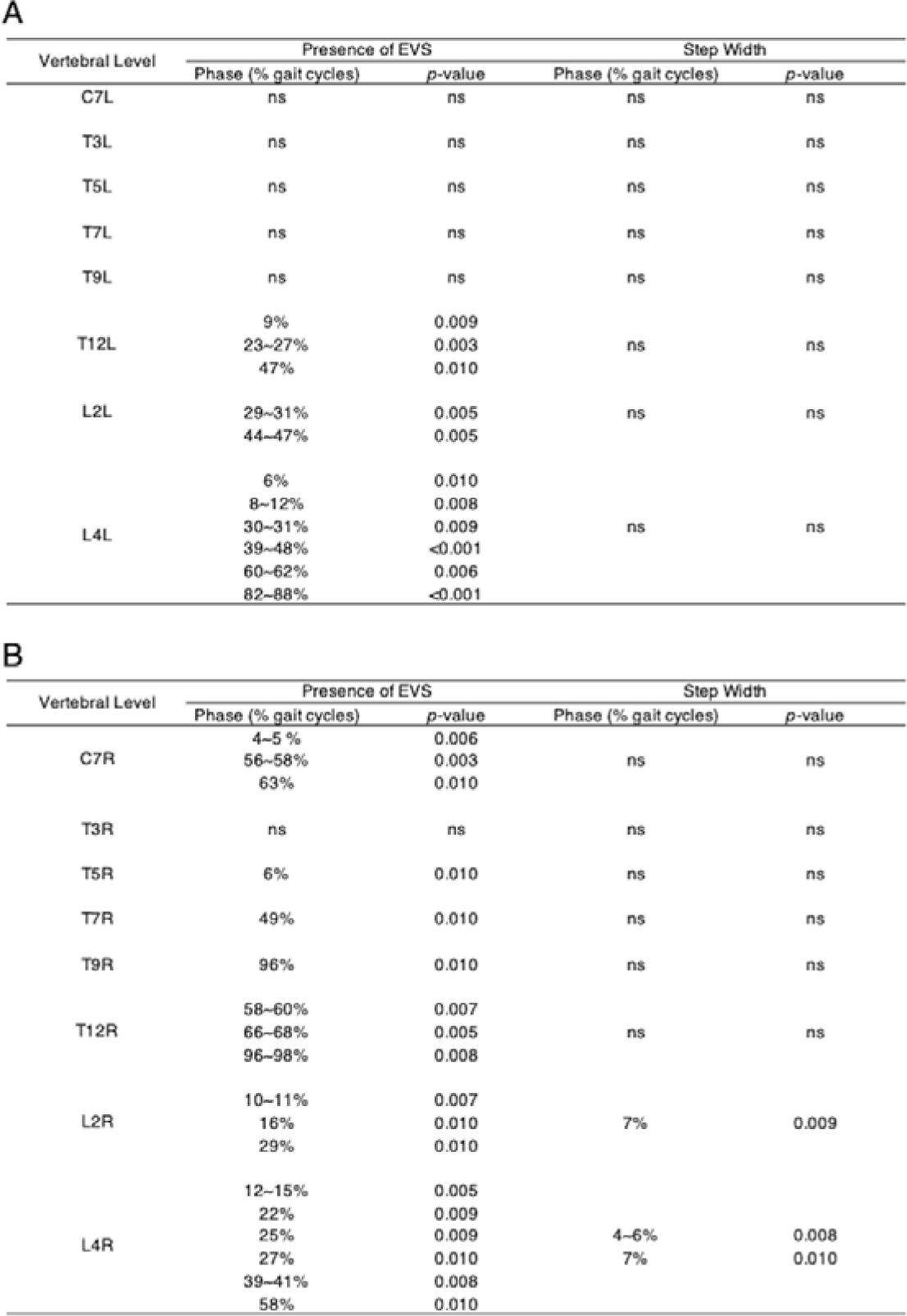
Summary of the effect of EVS and Step Width on Muscle Activity at Eight Vertebral Levels on the Left (A) and Right Side (B) Relative to the Midline of Trunk. Results from the One-dimensional Statistical Parametric Mapping based repeated-measures two-ways ANOVAs. The *p*-value indicates the probability that a suprathreshold cluster of the same spatiotemporal extent could have resulted from a random field process of the same smoothness as the observed residuals.

### EVS-EMG Coherence During Walking with Preferred Step Width

Significant EVS-EMG coupling, defined as coherence over 0.015, was observed in all recorded sites when walking with preferred step width. But differences between vertebral levels as to when, and in which frequency band this coherence occurred were found (Figure 5).

**Figure 5.**
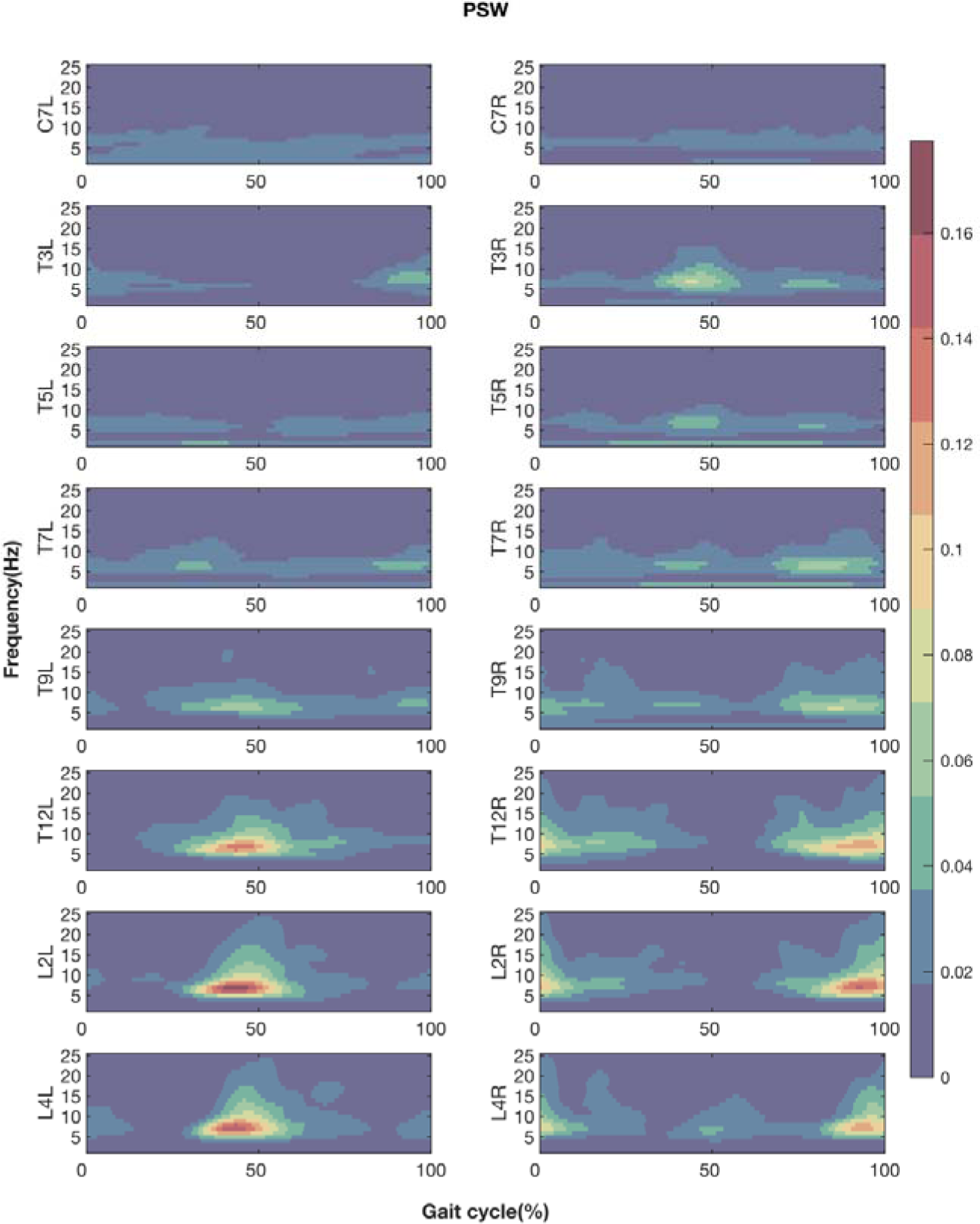
EVS-EMG Coherence Throughout the Gait Cycle During Walking with Preferred Step Width. Coherence between EVS and EMG of paraspinal muscles at C7, T3, T5, T7, T9, T12, L2 and L4 levels (from top to bottom) during walking with preferred step widths while exposed to EVS. The left and right columns represent the recorded sites at the left and right side relative to the mid-line of the trunk. The X-axis is the time (normalized to gait cycle, starting at the right heel strike). The Y-axis is the frequency range (from 1Hz to 25 Hz). Coherence amplitude is indicated by the colour bar. For illustrative purposes, only coherence above 0.015 are presented.

Phase-specific EVS-EMG coherence was roughly mirrored in the left and right relative to the midline of trunk, i.e., phase shifted by about half a gait cycle. Cluster-based permutation paired t-test showed no significant differences between coherence on the left and right side when they were shifted by half a gait cycle.

At vertebral levels from T12 to L4, peak coherence occurred around the ipsilateral heel strikes. At T7 and T9, peak coherence occurred before both ipsilateral and contralateral heel strikes. At vertebral levels from T3 to T5, peak coherence occurred around the contralateral heel strikes. At C7 level, coupling between EVS and EMG was almost continuously present, without distinct peaks, albeit at an overall lower magnitude. Overall, the phase-specific responses at the higher and lower vertebral levels were also mirrored, such that at lower levels a peak occurred at ipsilateral heel strike, while at higher levels, it occurred at contralateral heel strike.

Additionally, responses occurred earlier in the gait cycle at the higher vertebral levels compared to the lower levels. The magnitude of coherence was higher at the lower vertebral levels. Peak coherence aligned with one of the peaks in muscle activity at the lower vertebral levels (i.e., ipsilateral heel strike), and with the only peak, at contralateral heel strikes, for the higher vertebral levels. Moreover, coherence calculated from the last 50 gait cycles was on average lower than those calculated from the first 50 gait cycles. However, this difference was not significant.

### EVS-EMG Coherence During Walking with Narrow Step Width

To address our hypothesis that the magnitude of coupling between EVS and EMG during walking with narrow step width is larger than walking with preferred step width, the differences in EVS-EMG coherence between walking with preferred step width (PSW) and walking with narrow step width (NSW) were examined (Figure 6B).

**Figure 6.**
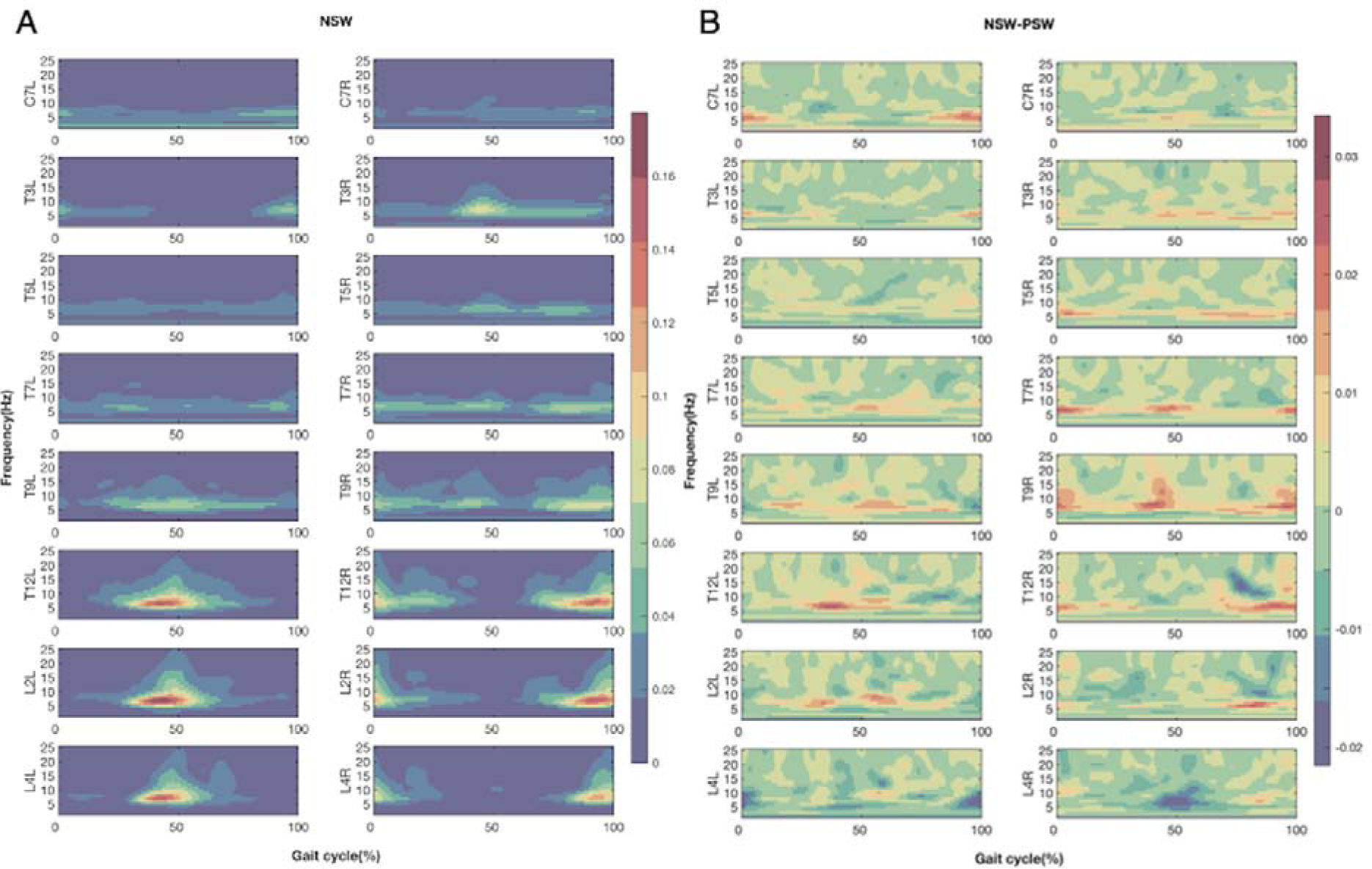
EVS-EMG Coherence in NSW and Effects of Step Width. (A) The coherence between EVS and EMG when walking with narrow step width (NSW); (B) The difference in coherence between walking with narrow step widths and preferred step width (NSW - PSW). Figures from top to bottom rows present the results at eight vertebral levels. The left or right columns represent the recorded sites at the left or right side relative to the mid-sagittal line of trunk. The X-axis is the time (normalized to gait cycle, starting at the right heel strike). The Y-axis is the frequency range (from 1Hz to 25 Hz). The magnitudes of coherence in NSW (A) and the difference of coherence between NSW and PSW (B) are indicated by the colour bar.

At the majority of vertebral levels, during most of the phases in the gait cycle, EVS-EMG coupling during walking with a narrow step width was slightly, but not significantly, higher than that during walking with preferred step width (Figure 6A). The greatest differences were observed at the instants around the peak coherence in normal walking. Interestingly, at the L4 level on both sides, there was a decrease in coherence when walking with a narrow step width, around the contralateral heel strikes, but again this was not significant.

Similar as when walking with preferred step width, a nonsignificant decreased coherence was found in the last 50 gait cycles compared to the first 50 gait cycles.

### Gain and Delay Estimates Between EVS and EMG

No significant effects on gains of either Vertebral level (F (1.127, 30.429) = 1.641, p = 0.201), Condition (F (0.161, 4.347) = 0.744, p = 0. 396), or their interaction (F (1.127, 30.429) = 1.736, p = 0. 147) were found (Figure 7A).

**Figure 7.**
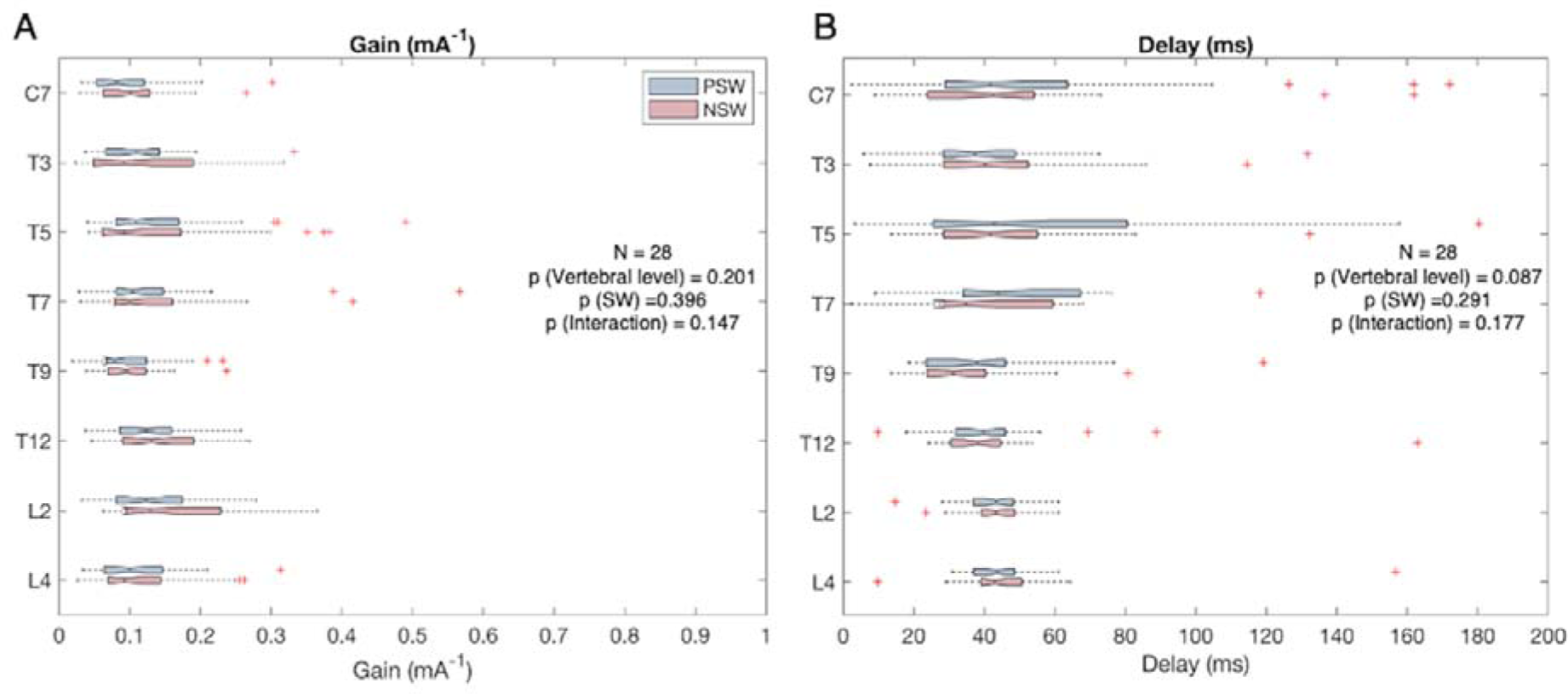
Gain(A) and Delay (B) Estimates Between EVS and EMG. The Y-axis represents vertebral levels (C7, T3, T5, T7, T9, T12, L2 and L4). In panel A, the X--1 axis represents the magnitude of the gain (mA) between EVS and EMG during walking with preferred (red box) and narrow step widths (blue box). In Panel B, the X-axis represents the delay (ms) between EVS and EMG. Within each box, the central mark indicates the median, and the bottom and top edges indicate the 25th and 75th percentiles, respectively. The red ‘+’ symbols represent the outliers, and the whiskers extend to the maximum and minimum values, excluding the outliers.

After Greenhouse-Geisser correction, no significant effects on delays of Vertebral level (F (2.415, 65.205) = 2.171, p = 0.087), Condition (F (0.161, 9.315) = 1.162, p = 0.291), or Vertebral level x Condition interaction (F (2.415, 65.205) = 1.638, p = 0.177) were found (Figure 7B).

## Discussion

We used electrical vestibular stimulation (EVS) and EMG to investigate the coupling of vestibular afference to paraspinal muscles activity at different vertebral levels. The Presence of EVS induced a significant increase in muscle activity at the lower vertebral levels around ipsilateral heel strikes, while the Step Width evoked a significant increase at the right L2 and L4 vertebral levels. EVS evoked significant increase in both step width and cadence. Variability of thorax roll significantly increased when walking with EVS with both step width conditions, while the variability of head rotation significantly decreased during walking with EVS with preferred step width. Significant coherence between the input EVS signal and output paraspinal muscle EMG, from cervical to lumbar vertebral levels, was found at ipsilateral and/or contralateral heel strikes. This coupling between EVS and EMG was roughly mirrored between the left and right body sides, i.e., phase shifted about half a gait cycle. Between the higher and lower vertebral levels, the coupling was also mirrored, such that a peak coherence occurred at ipsilateral heel strike at lower vertebral levels, while at higher levels it occurred at contralateral heel strike. EVS-EMG coupling only partially coincided with the peak muscle activity. Small but non-significant increases were found in EVS-EMG coupling during walking with narrow step widths. A non-significant reduction in EVS-EMG coherence in the last 50 gait cycles compared to the first 50 gait cycles was found for walking with preferred and narrow step widths. Below, we discuss how these findings advance our understanding of the contribution of the vestibular system to the control of paraspinal muscles.

### Attenuated EVS-EMG Coherence through Habituation

Non-significant attenuation in EVS-EMG coherence was found in both step width conditions. To ascertain that attenuation had no effects on our main findings, we compared the coherence in the two step width conditions in the first 50 gait cycles, which also showed no significant difference in EVS-EMG.

Attenuation in coherence over time is in line with the literature. For example, the gain between sinusoidal EVS and centre of pressure movement declined over a five-minutes standing trial (Balter *et al*., 2004). Hannan et al. reported that in standing, the gain between stochastic EVS to GRF decreased about 18% in the first 40 seconds, while over a one-hour walking trial, the gain decreased by roughly 38% and the coherence decreased from 0.04 to 0.02 (Hannan *et al*., 2021). Such a reduction in evoked responses has been attributed to habituation (Balter *et al*., 2004; Hannan *et al*., 2021). It is believed that the nervous system can down-weight the sensory cues offering irrelevant or erroneous information, and up-weight alternative sources of sensory information, known as ‘sensory reweighting’ (Balter *et al*., 2004; Hannan *et al*., 2021). As a result, the induced response to an EVS stimulus will decrease over time.

### The Contribution of The Vestibular System to Paraspinal Muscle Activity

In line with previous study, muscle activity at the lower vertebral levels significantly increased induced by both the presence of stimulation and the narrowing the step width (Guillaud *et al*., 2020; Magnani *et al*., 2021). Similar to previous studies, paraspinal muscles showed peak activity around both heel strikes at the lower vertebral levels (Ivanenko *et al*., 2004; de Seze *et al*., 2008; Guillaud *et al*., 2020; Magnani *et al*., 2021) and only a single peak at contralateral heel strikes at the higher vertebral levels (Ivanenko *et al*., 2006). However, at the higher spinal levels, de Seze et al (2008) reported a double-peaked pattern with a relatively smaller magnitude at ipsilateral heel strike. The difference in peak muscle activity at higher vertebral levels could be attributed to the lower walking speed in our study (2.8 km/h) as compared to the study of de Seze et al (2008) at 5 km/h, since the trunk muscle activity increases with walking speed (Callaghan *et al*., 1999; Anders *et al*., 2007).

At the higher vertebral levels, we found the EVS-EMG coupling partially aligned with the only peak in EMG in the gait cycle at contralateral heel strike. At the lower vertebral levels, we found that the peak EVS-EMG coupling partially aligned with the peak muscle activity at ipsilateral heel strike. At the contralateral heel strike, no significant EVS-EMG coupling was observed, in spite of the peak in muscle activity. This indicates that high motoneuron excitability does not necessarily cause strong effects of vestibular input. Other studies also reported that peak coherences do not always align with peak EMG for a range of muscles (Dakin *et al*., 2013; Forbes *et al*., 2017; Magnani *et al*., 2021). Partial alignment between EVS-EMG coupling and muscle activity was demonstrated in the Gastrocnemius muscle where the increase in coherence paralleled the increase in muscle activity but peaked slightly earlier in the gait cycle than the muscle activity (Blouin *et al*., 2011).

At the lumbar vertebral levels, the difference in EVS-EMG coupling between ipsilateral and contralateral heel strikes might be explained by the functional differences of the two peak activities at these phases of the gait cycle. At lumbar levels, the activations were described as an eccentric contraction at contralateral heel strikes, and a concentric contraction at the ipsilateral heel strikes (Shiavi, 1985; Anders *et al*., 2007). Our results shows that only the activity at ipsilateral heel strikes of the lumbar paraspinal muscles was affected by vestibular input, which may signify its role in stabilizing the trunk over the new stance leg.

### Vertebral Level Specific Response to EVS

In contrast to the expectation that, given a longer efferent pathway, the response delays would increase from the higher to lower vertebral levels, we found no significant differences in delays between the vertebral levels. This is consistent with a previous study addressing paraspinal muscles from T7 to L4 (Guillaud *et al*., 2020). However, we found differences in the phases, in which EVS-EMG coherence occurred between vertebral levels. This indicates that the modulation of paraspinal muscles activity based on vestibular afference is different between vertebral levels. This vertebral level specific modulation may reflect differences in vestibulospinal tracts transmitting the signals to motoneurons between vertebral levels (Wilson & Yoshida, 1969; Wilson *et al*., 1970; Wilson *et al*., 1995; Kasumacic *et al*., 2010). Using selective lesions in the brainstem and/or spinal cord in neonatal mice, Kasumacic et al (2010) showed that the vestibular inputs to motoneurons at the upper cervical levels primarily derive from the medial vestibulospinal tracts, while at lower cervical levels, there was a substantial contribution from the lateral vestibulospinal tract. This may potentially explain the constant EVS-EMG coupling at C7 level in our study, which was different from the distinctly phasic responses at the other vertebral levels. The continuous coupling at C7 may be related to head stabilization, an important function of the medial vestibulospinal tracts (Goldberg & Cullen, 2011). At the lumbar levels in mice, vestibular input solely came from the lateral vestibulospinal tracts (Kasumacic *et al*., 2010). At thoracic levels, early vestibular inputs originated from the lateral vestibulospinal tract, with later inputs potentially transmitted by midline-crossing motoneuron dendrites and/or excitatory commissural interneurons (Kasumacic *et al*., 2010). Transmission of vestibular signals by different tracts might be expected to cause a difference in response delays. However, the faster conduction velocity (90 m/sec) of the longer lateral-vestibulospinal tract and a slower conduction velocity (60 m/sec) of medial vestibulospinal tract (Peterson *et al*., 1978) may result in a similar delay and synchronized modulation of paraspinal muscle activity at different vertebral levels as a consequence.

Interestingly, we found that coherent responses at the upper and lower vertebral levels were mirrored, that is to say; at the lower levels, coherence was present around ipsilateral heel strike, while at the higher levels, it was present around contralateral heel strike. From a biomechanical point of view, this spatial pattern of vestibular induced activation could ensure maintenance of head orientation in space, while moving the trunk to correct a deviation of the centre of mass. To be more specific, a left displacement of the centre of mass could be counteracted by the contraction of left lumbar muscles. To maintain the head orientation in space, cervical and upper thorax muscles activity at the right side would be needed. These mirrored responses between upper-ipsilateral and lower-contralateral muscles to the vestibular signals have also been observed in mice (Kasumacic *et al*., 2010). After ipsilateral hemi-sections of the spinal cord at C1 level in mice, the responses to electrical stimulation in vestibulospinal motoneurons at the contralateral side were completely eliminated at the thoracic and lumbar levels, while at the cervical level the contralateral responses remained unaffected, but disappeared after contralateral hemi-section, and vice versa (Kasumacic *et al*., 2010). Furthermore, signals from the ipsilateral-descending lateral vestibulospinal tract are transmitted to contralateral motoneurons by descending commissural interneurons. These interneurons are more prevalent at the T7 level than at L1, L2 and L5 level (Kasumacic *et al*., 2010; Kasumacic *et al*., 2015). This finding is in line with our observation that at T5 and T7 levels, the spatial pattern of EVS-EMG coupling resembles a combination of those at higher thoracic and lumbar levels.

### The Contribution of The Vestibular System to Gait and Postural Control

In line with a previous study, both step width and cadence significantly increased when walking with EVS in both step width conditions (Magnani *et al*., 2023). As reported previously, in response to perturbations, participants decrease their step length and increase their step width and cadence to maintain balance in the mediolateral direction (Hak *et al*., 2012).

The presence of EVS evoked opposite responses in head and thorax rotation variability, i.e., when walking with EVS with preferred step width, head rotation variability significantly decreased while thorax rotation variability significantly increased. These results are in line with an earlier study reporting increased averaged thorax rotation and decreased head rotation in roll during walking evoked by electrical vestibular stimulation (Deshpande *et al*., 2015). A possible reason for this might be that lumbar movements are elicited by EVS to counteract illusory movements while during walking the head is stabilized in space. To do so, a counter rotation at the thoracic or cervical levels is required. This is in line with the continuous EVS-EMG coupling at the C7 level and the mirrored EVS-EMG coupling between the thoracic and lumbar levels.

With restricted foot placement, more use of alternative strategies for balance control can be expected, i.e., larger shifts of the centre of pressure under the foot through ankle moments or larger changes in the body’s angular momentum through segmental movements relative to the body’s centre of mass (Hof, 2007).The latter could be expected to involve increased trunk and head roll movement. However, we found overall less thorax roll variability in the narrow than preferred step width conditions. This suggests that the contribution of upper body rotations to balance control is limited during narrow base walking. With the relatively narrow base of support provided by the narrow step width, trunk rotations might lead to instability. Thus, controlling the centre of mass by shifting the centre of pressure through ankle moments might be safer. This is in line with previous work showing that if foot placement cannot be used for balance control due to foot placement constraints, ankle moments are primarily used instead of angular momentum changes (van Leeuwen et al., 2022).

### Impact of Step Width on Vestibular Contributions

Surprisingly, our outcomes showed no statistically significant differences in EVS-EMG coherence between walking at preferred and narrow step widths. In contrast, in an earlier study, EVS-EMG coherence of the lumbar erector spinae muscle significantly increased when walking with narrower step widths, while in medial gastrocnemius and gluteus medius coherence decreased or remained unchanged, respectively (Magnani *et al*., 2021). However, we confirmed the hypothesis that EVS-evoked responses increase when walking with narrow step widths by observing a significant increase in the coupling between EVS and the horizontal ground reaction force in medial-lateral direction during mid-stance phase (Figure 9). This observation is consistent with Magnani et al. (2021).

The possible reasons of finding the opposite in EVS-EMG coherence can be: 1) the ground reaction force is the summation of all muscle activity, it is reasonable to find the significant difference in EVS-GRF coherence but not in paraspinal muscle EVS-EMG coherence; 2) participants did not fully keep to our instructions of halving step width; 3) we did not ask participants to walk with an upward head tilt of 18 degree as Magnani et al. (2021). Such a posture maximizes the perception of head roll during EVS (Fitzpatrick & Day, 2004; St George & Fitzpatrick, 2011; Magnani *et al*., 2021). Since we imposed step width by lines projected on the treadmill, it was impossible to keep this head orientation. Besides, this non neutral posture can lead to a different trunk posture resulting in differences in trunk muscle activity. Moreover, the tilted head posture can limit visual cues. Based on the theory of sensory reweighting, vestibular information might have been up-weighted amplifying any effects on EVS-EMG coherence.

**Figure 8.**
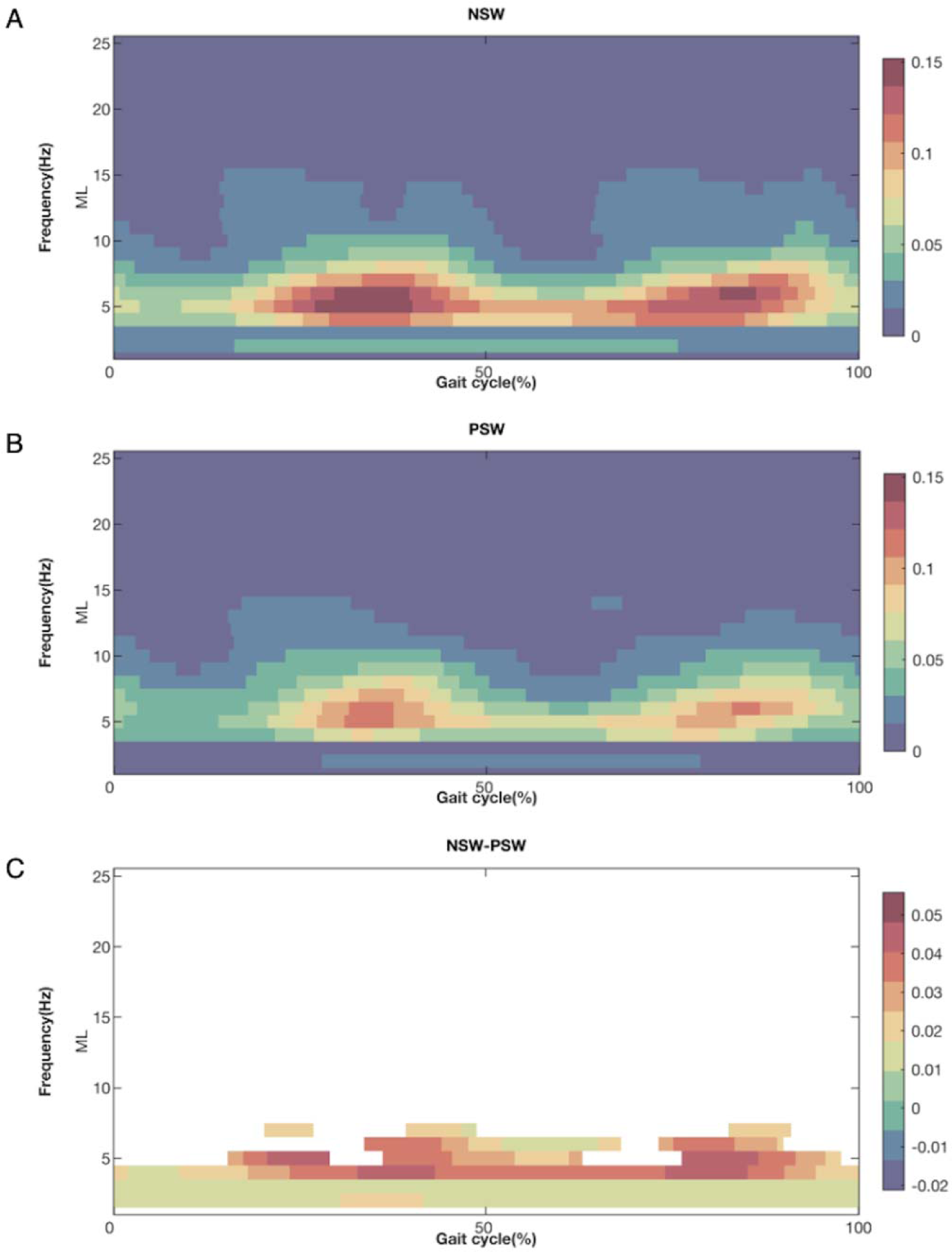
EVS-GRF Coherence in NSW (A), PSW (B) and Effects of Step Width (C). The coherence between EVS and GRF in medial-lateral (ML) when walking with narrow step width (NSW) (A) and preferred step width (PSW) (B); (C) The difference in coherence between walking with narrow step widths and preferred step width (NSW - PSW). The X-axis is the time, normalized to gait cycle, starting at the right heel strike. The Y-axis is the frequency range (from 1Hz to 25 Hz). The magnitudes of coherence in NSW (A), PSW (B) and the difference of coherence between NSW and PSW (C) are indicated by the colour bar. For illustrative purposes, only significant difference was plotted.

### Limitations

One important limitation of this study is that currently there is no validated method to eliminate the artifacts of electrical stimulation from EMG signals. Although the filtered EMG still contains the timing information of activation in muscles evoked by stimulation (Forbes *et al*., 2018), it is unclear how appropriate the filtering is. Filtering EMG data to remove the EVS artifacts will inevitably also remove EMG and while information on lower bandwidth changes in muscle force is largely maintained or even enhanced (Potvin & Brown, 2004), information on muscle activation in the frequency range of interest here may be lost due this procedure. In addition, rectification of the EMG signals may cause a nonlinear distortion (Neto & Christou, 2010; McClelland *et al*., 2014), which may enhance (Ward *et al*., 2013) or decrease coherence (Neto & Christou, 2010) or create false coherence in a frequency range where there is no input (Neto & Christou, 2010). Still, the application of filtering and rectification is reasonable in view of our primary interest in the modulation of muscle activation and coherence over the gait cycle (Myers *et al*., 2003; Halliday & Farmer, 2010; van Asseldonk *et al*., 2014; Jiang *et al*., 2019). Nevertheless, finding an appropriate method for artifact removal may further strengthen our understanding of the electrical stimulus-response relationship.

### Conclusion

We explored how vestibular signals influence paraspinal muscle activity at different vertebral levels in walking with preferred and narrow step width. EVS and the modulation of step with induced significant increase in muscle activity at the lower vertebral levels. Significant EVS-EMG coupling was observed at all recorded sites around the ipsilateral and/or contralateral heel strikes. This coupling was vertebral level specific and exhibited a mirrored pattern between the left and right side of the body, as well as between the higher and lower vertebral levels. In other words, at the same vertebral level, the coupling on the left side exhibited a phase shift of half a gait cycle compared to the right side. Along the vertebral levels, a peak occurred at ipsilateral heel strike at lower levels, while it occurred at the contralateral heel strike at higher levels. We found that the EVS-EMG coupling partially coincided with the peak muscle activity. No significant differences were found in gain and delay between either the vertebral levels or step width conditions. Significant increase was found in cadence and step width when walking with EVS in two step width conditions. EVS evoked opposite responses in head and thorax rotation, i.e., the head rotation variability significantly decreased while the thorax rotation variability significantly increased when walking with EVS with preferred step width. Vertebral level specific modulation of paraspinal muscle activity based on vestibular signals might allow a fast, synchronized, and spatially coordinated response along the trunk during walking.

## Additional information

### Data Availability Statement

The original data are available from the corresponding author upon reasonable request.

### Competing Interests

The authors declare that there are no conflicts of interest.

### Author Contributions

Experiments were performed at the Dual Belt Lab in the Vrije Universiteit Amsterdam. Conception and design: Y.C.L., S.M.B., K.K.L. and J.H.D. Data acquisition: Y.C.L. Analysis and interpretation: Y.C.L., S.M.B., K.K.L., S.B. and J.H.D. Drafting and revising manuscript: Y.C.L., S.M.B., K.K.L., S.B. and J.H.D. All authors have read and approved the final version of this manuscript and agree to be accountable for all aspects of the work in ensuring that questions related to the accuracy or integrity of any part of the work are appropriately investigated and resolved. All persons designated as authors qualify for authorship, and all those who qualify for authorship are listed.

### Funding

Y.C.L. was funded by a scholarship (No. 202108520034) from the China Scholarship Council (CSC). S.M.B. was funded by a VIDI grant (no. 016.Vidi.178.014) from the Dutch Organization for Scientific Research (NWO).

## Acknowledgements

The authors appreciate all participants for their voluntary participation. The authors gratefully acknowledge Dr. Patrick A. Forbes from Erasmus MC for sharing the MATLAB scripts to calculate the coherence and frequency response function. The authors would like to thank Dr. Mohammadreza Mahaki for his help in data collection. The authors thank Bert Clairbois and Richard Casius for the technical assistance.

## Notes

### Competing Interest Statement

The authors have declared no competing interest.

### Summary of Updates

In Result, section 'Effects of EVS and Step Width on Gait Parameters' and 'Effects of EVS and Step Width on Posture Control ' was added to investigate the effect of vestibular stimulation on controlling the gait and posture. In Discussion, we added 'Attenuated EVS-EMG Coherence through Habituation', 'The Contribution of The Vestibular System to Gait and Postural Control ' and 'Figure 8 EVS-GRF Coherence in NSW (A), PSW (B) and Effects of Step Width (C)' to discuss revised results.

